# Disparity, Diversity, and Duplications in the Caryophyllales

**DOI:** 10.1101/132878

**Authors:** Stephen A. Smith, Joseph W. Brown, Ya Yang, Riva Bruenn, Chloe P. Drummond, Samuel F. Brockington, Joseph F. Walker, Noah Last, Norman A. Douglas, Michael J. Moore

## Abstract

- The role whole genome duplication (WGD) plays in the history of lineages is actively debated. WGDs have been associated with advantages including superior colonization, various adaptations, and increased effective population size. However, the lack of a comprehensive mapping of WGDs within a major plant clade has led to uncertainty regarding the potential association of WGDs and higher diversification rates.
- Using seven chloroplast and nuclear ribosomal genes, we constructed a phylogeny of 5,036 species of Caryophyllales, representing nearly half of the extant species. We phylogenetically mapped putative WGDs as identified from analyses on transcriptomic and genomic data and analyzed these in conjunction with shifts in climatic niche and lineage diversification rate.
- Thirteen putative WGDs and twenty-seven diversification shifts could be mapped onto the phylogeny. Of these, four WGDs were concurrent with diversification shifts, with other diversification shifts occurring at more recent nodes than WGDs. Five WGDs were associated with shifts to colder climatic niches.
- While we find that many diversification shifts occur after WGDs it is difficult to consider diversification and duplication to be tightly correlated. Our findings suggest that duplications may often occur along with shifts in either diversification rate, climatic niche, or rate of evolution.

## Introduction

Understanding the causes and correlates of diversification within flowering plants has been a central goal of evolutionary biologists. Genomic and transcriptomic data have reinvigorated hypotheses associating whole genome duplication (WGD) with lineage diversification rate increases (e.g., Levin, 1983; Levin 2002; Barker *et al*. 2009; Estep *et al.*, 2014; Soltis *et al*., 2014; Edger *et al*. 2015; Puttick *et al*. 2015; Tank *et al*., 2015; Barker *et al*. 2016; Huang *et al*. 2016; McKain *et al*. 2016; Laurent *et al*. 2017). It is not self-evident why WGDs would be associated with increases in lineage diversification. One hypothesis suggests that the additional genetic material provides a basis to generate new adaptations (Edger *et al*., 2015), although this itself assumes a co-occurrence of adaptation and lineage proliferation (Levin, 1983). The apparent lack of precise co-occurrence of adaptation and lineage proliferation has been explained by the potential of a lag model (Schranz *et al*. 2012; Tank *et al*. 2015) where diversification may follow WGD events. In the absence of overwhelming correlative signal, we are often unable to discern true ancient WGD events from aneuploidy without advanced genomic information such as synteny mapping (Dohm *et al*., 2012). Because it is often difficult to distinguish the two, for simplicity we will define WGD broadly to include putative ancient WGD events (paleopolyploidy) and ancient aneuploidy events. WGD events are thought to be a common occurrence and have been associated with an estimated 15% of angiosperm speciation events (Wood *et al*., 2009). However, whether speciation by WGD is correlated with higher diversification rates remains highly debated (Mayrose *et al*., 2011; Estep *et al*. 2014; Soltis *et al*., 2014; Tank *et al*., 2015; Kellogg *et al*. 2016). Analyses based on recent WGD events have concluded that immediate extinction rates are higher for polyploid plants (Mayrose *et al*., 2011; Arrigo and Barker, 2012). This may result from small initial population sizes and an increased dependence on selfing. Alternatively, despite the disadvantages of WGD, others have suggested that polyploids may be superior colonizers (Soltis and Soltis, 2000).

Indeed, extreme environments are associated with high levels of WGD, with up to 87% of species restricted to areas that were glaciated during the last ice age consisting of polyploids (Brochmann, 2004). However, in the example from Arctic plants, the high level of WGD has occurred post-glaciation representing a micro-evolutionary period whereas previous studies often focus at much deeper macro-evolutionary time scales (Mayrose *et al*., 2011; Tank *et al*., 2015; Soltis *et al*., 2014). From the perspective of a short timescale, polyploidy has the disadvantages of higher error rates in mitosis (Storchová *et al*., 2006) and masking of deleterious mutations allowing them to accumulate to higher frequencies in a population (Otto & Whitton, 2000). A suite of advantages however may also arise, including gain of asexuality (Miller *et al*. 2000) and varying effects of heterosis (Comai, 2005). The net role these advantages and disadvantages play on the macroevolutionary scale is difficult to determine from either the purely short-term or purely long-term time scales previously used.

The long-term consequence of WGD is a central question in macroevolution and comparative genomics. However, with a suite of advantages and disadvantages, much debate surrounds the importance and patterns of correlation of WGD (Comai 2005). While polyploidization events can cause instant speciation, there is no reason to assume that these singular speciation events in themselves would influence large-scale diversification rate shifts when considering lineage survivorship. Instead, there may be other factors, such as the increase in genetic material, perhaps increasing genetic diversity or enabling adaptation, that cause long term shifts in rates of diversification. Adaptations need not be associated with shifts in the tempo of diversification and those adaptations and shifts in diversification may not co-occur on the same branch (i.e., there may be a lag time; Donoghue 2005; Smith *et al.*, 2011, Schranz *et al.*, 2012; Donoghue & Sanderson 2015; Tank *et al.*, 2015; Dodsworth *et al.*, 2016). In the broader context of plant evolution, there are several possible outcomes of WGD in relation to the evolution and diversification of clades: 1) no relationship between WGD and speciation rate or habitat shift/adaptation, 2) WGD coincides with an increase of speciation rate, with or without a lag time, 3) WGD promotes dispersal and habitat shifts, which has mixed relationship with speciation rate, and 4) a mixture (some association, some not), similar to the previous hypothesis but without explicitly promoting dispersal or habitat shift or speciation (e.g., adaptation could be more prominent than dispersal and habitat shift). Here, we contribute to this discussion on diversification and WGDs with an in-depth examination of the intersection of diversification and WGDs happening at a range of scales within the hyperdiverse Caryophyllales.

The Caryophyllales comprise ~12,500 species in 39 families (Thulin *et al*., 2016; APG IV: Chase *et al*., 2016), representing approximately 6% of extant angiosperm species diversity. The estimated crown age of Caryophyllales is approximately 67–121 millions of years ago (mega-annum, Ma) (Bell *et al*., 2010; Moore *et al*., 2010). Species of the Caryophyllales exhibit extreme life-history diversity, ranging from tropical trees to temperate annual herbs, and from desert succulents (e.g., Cactaceae) to a diverse array of carnivorous plants (e.g., the sundews *Drosera* and pitcher plants *Nepenthes*). Such extraordinary diversity makes Caryophyllales a particularly useful system for investigating the relationship between WGD vs. diversification and niche evolution. Our previous analyses using 62 transcriptomes representing 60 species across the Caryophyllales identified 13 well-supported ancient WGD events (Yang *et al*., 2015). We have since nearly tripled the taxon sampling and assembled a data set comprising high-coverage transcriptomes and genomes from 169 species across the Caryophyllales (Yang *et al*., submitted), providing even greater power for resolving the number and phylogenetic locations of WGD events. Moreover, the growth in the number of plant taxa on GenBank that are represented by traditional targeted sequences (e.g., *rbc*L, *mat*K, ITS, etc.) and the growth of publicly available collections data (e.g., GBIF, iDigBio) provide excellent opportunities to apply megaphylogeny and niche diversification approaches at fine scales in Caryophyllales.

By examining WGDs and diversification within the Caryophyllales, we present an important example. Not only does the dataset examined have a high density of transcriptomic sampling, the diversification of the bulk of Caryophyllales occurred during a time frame intermediate to that of most published studies that have probed a link between WGD and macroevolution. This time frame, between 10 and 100 Ma, is important for angiosperms as much of the diversification that has led to the modern flora occurred during this period and most modern angiosperm families appeared by this time. Discussion of speciation rate, niche shift, and WGD would be flawed without accurate mappings of WGD events within this time scale. We compiled a data set with extensive and precise mapping of WGD combined with a species-level phylogeny. The megaphylogeny approach has been used extensively in the past to combine data from many gene regions and across broad taxonomic groups to address evolutionary questions (Smith *et al*., 2009). Here, we use this approach to help inform analyses from phylogenomic studies, and provide a broad context in which to examine these genomic phenomena. With half of the species sampled, this represents one of the largest and most exhaustive studies of WGDs, diversification rate, and adaptive shifts.

## Materials and Methods

### Sanger sequencing and assembly

A total of 248 new *matK* sequences were included in this study (Table 1). To generate these sequences, leaf samples were collected in silica in the field or from cultivated material, or were collected from herbarium sheets. DNA was isolated using either the Nucleon Phytopure kit (GE Healthcare Life Sciences, Pittsburgh, PA, USA), using the 0.1 g protocol and following manufacturer’s instructions, or using the Doyle & Doyle (1987) protocol, with the addition of 1% PVP-40. An approximately 950 bp region in the middle of the *matK* gene was amplified and sequenced using custom-designed primers (Table 2). PCRs were performed in 12.5 µL volumes with 0.5 µL of 5 mM primer for both primers, 5-20 ng of DNA template, 0.1 µL of GoTaq (Promega, Madison, WI, USA), 6.25 µL of Failsafe Premix B (Epicentre, Madison, WI, USA), and 4.7 µL of sterile, deionized water. Reactions were run on a Bio-Rad PTC 200 thermocycler (Bio-Rad, Hercules, CA, USA) at Oberlin College. Individual PCRs were cleaned in 16.5 µL reactions containing 10 U of Exonuclease I (Affymetrix, ThermoFisher Scientific, Waltham, MA, USA), 2 U of shrimp alkaline phosphatase (Affymetrix), 8 µL of PCR product, and 8.5 µL of sterile, deionized water. Sanger sequencing of the resulting cleaned PCRs was conducted by Neogenomics (formerly SeqWright; Houston, TX, USA) using an ABI 3730xl automated sequencer (Applied Biosystems, ThermoFisher Scientific). The resulting forward and reverse sequences for each reaction were trimmed and *de novo* assembled using default parameters of the Geneious assembler in Geneious versions 5-7 (Biomatters, Auckland, New Zealand).

**Table 1:** Voucher information and GenBank accession numbers for newly reported plastid *matK* sequences. Families follow APG IV (Angiosperm Phylogeny Group, 2016).

| Family | Taxon | Voucher specimen<br>(Herbarium acronym) | Collection<br>locality | NCBI<br>accession<br>number |
| --- | --- | --- | --- | --- |
| Achatocarpaceae | *Achatocarpus gracilis*<br>H.Walter | Silvia H. Salas<br>Morales et al. 5608<br>(TEX) | Mexico:<br>Oaxaca | KY952292 |
| Achatocarpaceae | *Phaulothamnus*<br>*spinescens* A.Gray | Michael J. Moore et<br>al. 976 (OC) | United States:<br>Texas | KY952477 |
| Achatocarpaceae | *Phaulothamnus*<br>*spinescens* A.Gray | William R. Carr<br>27176 (TEX) | United States:<br>Texas | KY952478 |
| Amaranthaceae | *Allenrolfea occidentalis*<br>(S.Watson) Kuntze | Michael J. Moore 474<br>(OC) | United States:<br>Texas | KY952314 |
| Amaranthaceae | *Alternanthera*<br>*caracasana* Kunth | Michael J. Moore<br>1808 (OC) | United States:<br>Texas | KY952319 |
| Amaranthaceae | *Amaranthus cruentus* L. | Michael J. Moore 356<br>(OC) | United States:<br>Ohio<br>(cultivated) | KY952320 |
| Amaranthaceae | *Amaranthus* sp. | Michael J. Moore<br>1801 (OC) | United States:<br>Texas | KY952321 |
| Amaranthaceae | *Amaranthus* sp. | Michael J. Moore<br>2186 (OC) | United States:<br>Ohio | KY952322 |
| Amaranthaceae | *Amaranthus* sp. | Michael J. Moore<br>2187 (OC) | United States:<br>Illinois | KY952323 |
| Amaranthaceae | *Atriplex prosopidum*<br>I.M.Johnst. | Hilda Flores Olvera<br>et al. 1658 (MEXU) | Mexico:<br>Coahuila | KY952340 |
| Amaranthaceae | *Atriplex* sp. | Michael J. Moore<br>1689 (OC) | United States:<br>Texas | KY952338 |
| Amaranthaceae | *Atriplex* sp. | Michael J. Moore<br>1699 (OC) | United States:<br>Texas | KY952339 |
| Amaranthaceae | *Celosia argentea* L. var.<br>*plumosa* | Michael J. Moore 359<br>(OC) | United States:<br>Ohio<br>(cultivated) | KY952359 |
| Amaranthaceae | *Charpentiera ovata*<br>Gaudich. var. *ovata* | Flora K. Samis 7<br>(Lyon Arboretum<br>living collection,<br>accession 2011.0034) | United States:<br>Hawaii | KY952360 |
| Amaranthaceae | *Charpentiera tomentosa*<br>Sohmer var.<br>*maakuaensis* Sohmer | Flora K. Samis 6<br>(Lyon Arboretum<br>living collection,<br>accession 88.0141) | United States:<br>Hawaii | KY952361 |
| Amaranthaceae | *Chenopodium album* L. | Michael J. Moore 344<br>(OC) | United States:<br>Ohio | KY952362 |
| Amaranthaceae | *Gossypianthus*<br>*lanuginosus* (Poir.) Moq. | Michael J. Moore<br>1807 (OC) | United States:<br>Texas | KY952408 |
| Amaranthaceae | *Guilleminea densa* | Michael J. Moore et | Mexico: | KY952412 |
|  | (Humb. & Bonpl. ex Schult.) Moq. | al. 2445 (OC) | Chihuahua |  |
| Amaranthaceae | *Kali tragus* (L.) Scop. | Michael J. Moore 453 (OC) | United States: Texas | KY952506 |
| Amaranthaceae | *Nototrichium divaricatum* D.H.Lorence | Flora K. Samis 3 (Lyon Arboretum living collection, accession 96.0036 #3) | United States: Hawaii | KY952468 |
| Amaranthaceae | *Nototrichium humile* Hillebr. | Flora K. Samis 2 (Lyon Arboretum living collection, accession 2001-0254) | United States: Hawaii | KY952469 |
| Amaranthaceae | *Suaeda jacoensis* I.M.Johnst. | Hilda Flores Olvera et al. 1662 (MEXU) | Mexico: Coahuila | KY952514 |
| Amaranthaceae | *Suaeda jacoensis* I.M.Johnst. | Michael J. Moore et al. 2617 (OC) | Mexico: Nuevo Leon | KY952515 |
| Amaranthaceae | *Suaeda mexicana* (Standl.) Standl. | Hilda Flores Olvera et al. 1654 (MEXU) | Mexico: Coahuila | KY952516 |
| Amaranthaceae | *Tidestromia lanuginosa* (Nutt.) Standl. | Michael J. Moore 1128 (OC) | United States: Texas | KY952521 |
| Amaranthaceae | *Zuckia brandegeei* (A.Gray) S.L.Welsh & Stutz var. *plummeri* (Stutz & S.C.Sand.) Dorn | Joseph L. M. Charboneau 9672 (RM) | United States: Colorado | KY952528 |
| Cactaceae | *Leuenbergeria quisqueyana* (Alain) Lodé | Flora K. Samis 11 (Lyon Arboretum living collection, accession 2000.0281) | United States: Hawaii | KY952473 |
| Caryophyllaceae | *Moehringia macrophylla* (Hook.) Fenzl | Arianna Goodman 1 (OC) | United States: Oregon | KY952464 |
| Caryophyllaceae | *Paronychia lundellorum* Torr. & A.Gray | William R. Carr 17607 (MEXU) | United States: Texas | KY952472 |
| Caryophyllaceae | *Saponaria officinalis* L. | Michael J. Moore et al. 1819 (OC) | United States: Indiana | KY952507 |
| Caryophyllaceae | *Schiedea kaalae* Wawra | Flora K. Samis 5 (Lyon Arboretum living collection, accession 92.0513) | United States: Hawaii | KY952509 |
| Caryophyllaceae | *Spergularia salina* J.Presl & C.Presl | Michael J. Moore 1693 (OC) | United States: Texas | KY952512 |
| Didiereaceae | *Alluaudia ascendens* (Drake) Drake | Michael J. Moore 1645 | United States (cultivated) | KY952318 |
| Dioncophyllaceae | *Triphyophyllum peltatum* (Hutch. & Dalziel) Airy Shaw | Carel C. H. Jongkind et al. 7136 (WAG) | Liberia | KY952524 |
| Droseraceae | *Drosera burmannii* Vahl cv. Pilliga Red | Michael J. Moore 1814 (OC) | United States (cultivated) | KY952400 |
| Droseraceae | *Drosera peltata* Thunb. | Michael J. Moore | Australia: | KY952401 |
|  |  | 1817 (OC) | Tasmania<br>(cultivated) |  |
| Droseraceae | *Drosera regia* Stephens | Michael J. Moore<br>1812 (OC) | United States<br>(cultivated) | KY952402 |
| Drosophyllaceae | *Drosophyllum*<br>*lusitanicum* (L.) Link | Michael J. Moore<br>1816 (OC) | United States<br>(cultivated) | KY952403 |
| Frankeniaceae | *Frankenia gypsophila*<br>I.M.Johnst. | Michael J. Moore et<br>al. 1880 (OC) | Mexico:<br>Nuevo Leon | KY952406 |
| Microteaceae | *Microtea debilis* Sw. | Manuel Rimachi<br>11128 (TEX) | Peru: Loreto | KY952415 |
| Montiaceae | *Claytonia sibirica* L. | Arianna Goodman 2<br>(OC) | United States:<br>Oregon | KY952363 |
| Montiaceae | *Phemeranthus*<br>*parviflorus* (Nutt.) Kiger | Michael J. Moore et<br>al. 2214 (OC) | United States:<br>New Mexico | KY952479 |
| Nyctaginaceae | *Abronia angustifolia*<br>Greene | Michael J. Moore et<br>al. 2063 (OC) | Mexico:<br>Coahuila | KY952281 |
| Nyctaginaceae | *Abronia angustifolia*<br>Greene | Michael J. Moore et<br>al. 896 (OC) | United States:<br>New Mexico | KY952282 |
| Nyctaginaceae | *Abronia bigelovii*<br>Heimerl | Michael J. Moore et<br>al. 704 (OC) | United States:<br>New Mexico | KY952283 |
| Nyctaginaceae | *Abronia elliptica*<br>A.Nelson | Norman A. Douglas<br>2039 (DUKE) | United States:<br>Arizona | KY952284 |
| Nyctaginaceae | *Abronia fragrans* Nutt.<br>ex Hook. | Billie L. Turner 20-22 (SRSC) | United States:<br>Texas | KY952285 |
| Nyctaginaceae | *Abronia fragrans* Nutt.<br>ex Hook. | Glenn Kroh et al.<br>3021 (TEX) | United States:<br>Texas | KY952286 |
| Nyctaginaceae | *Abronia macrocarpa*<br>L.A.Galloway | Steve L. Orzell et al.<br>6492 (TEX) | United States:<br>Texas | KY952287 |
| Nyctaginaceae | *Abronia mellifera*<br>Douglas ex Hook. | N. Elizabeth<br>Saunders BP 19<br>(SIU) | United States:<br>Wyoming | KY952288 |
| Nyctaginaceae | *Abronia mellifera*<br>Douglas ex Hook. | N. Elizabeth<br>Saunders BP 20<br>(SIU) | United States:<br>Wyoming | KY952289 |
| Nyctaginaceae | *Abronia nana* S.Watson<br>var. *nana* | Robert C. Sivinski et<br>al. 3108 (NMC) | United States:<br>Arizona | KY952290 |
| Nyctaginaceae | *Abronia umbellata* Lam. | N. Elizabeth<br>Saunders LU 45<br>(SIU) | United States:<br>California | KY952291 |
| Nyctaginaceae | *Acleisanthes acutifolia*<br>Standl. | James Henrickson et<br>al. 22916 (TEX) | Mexico:<br>Coahuila | KY952293 |
| Nyctaginaceae | *Acleisanthes*<br>*angustifolia* (Torr.)<br>R.A.Levin | Michael J. Moore 460<br>(OC) | United States:<br>Texas | KY952294 |
| Nyctaginaceae | *Acleisanthes* cf.<br>*purpusiana* (Heimerl)<br>R.A.Levin | James Henrickson<br>23026 (TEX) | Mexico:<br>Coahuila | KY952309 |
| Nyctaginaceae | *Acleisanthes*<br>*chenopodioides*<br>(A.Gray) R.A.Levin | Michael J. Moore et<br>al. 733 (OC) | United States:<br>Texas | KY952295 |
| Nyctaginaceae | *Acleisanthes crassifolia*<br>A.Gray | Michael J. Moore et al. 569 (OC) | United States: Texas | KY952296 |
| Nyctaginaceae | *Acleisanthes diffusa*<br>(A.Gray) R.A.Levin var. *diffusa* | Michael J. Moore et al. 624 (OC) | United States: Texas | KY952297 |
| Nyctaginaceae | *Acleisanthes lanceolata*<br>(Wooton) R.A.Levin var. *lanceolata* | Michael J. Moore et al. 870 (OC) | United States: New Mexico | KY952298 |
| Nyctaginaceae | *Acleisanthes lanceolata*<br>(Wooton) R.A.Levin var. *lanceolata* | Michael J. Moore et al. 903 (OC) | United States: Texas | KY952299 |
| Nyctaginaceae | *Acleisanthes lanceolata*<br>(Wooton) R.A.Levin var. *megaphylla*<br>(B.A.Fowler & B.L.Turner) Spellb. & J.Poole | Alfred T. Richardson 1666 (TEX) | Mexico: Chihuahua | KY952300 |
| Nyctaginaceae | *Acleisanthes longiflora*<br>A.Gray | Michael J. Moore 435 (OC) | United States: Texas | KY952301 |
| Nyctaginaceae | *Acleisanthes longiflora*<br>A.Gray | Michael J. Moore et al. 571 (OC) | United States: Texas | KY952302 |
| Nyctaginaceae | *Acleisanthes nana*<br>I.M.Johnst. | Jackie Smith et al. 798 (TEX) | Mexico: San Luis Potosi | KY952303 |
| Nyctaginaceae | *Acleisanthes obtusa*<br>(Choisy) Standl. | Michael J. Moore et al. 984 (OC) | United States: Texas | KY952304 |
| Nyctaginaceae | *Acleisanthes palmeri*<br>(Hemsley) R.A.Levin | George S. Hinton 28620 (TEX) | Mexico: Nuevo Leon | KY952305 |
| Nyctaginaceae | *Acleisanthes parvifolia*<br>(Torr.) R.A.Levin | Michael J. Moore 452 (OC) | United States: Texas | KY952306 |
| Nyctaginaceae | *Acleisanthes purpusiana*<br>(Heimerl) R.A.Levin | James Henrickson 22709 (TEX) | Mexico: Coahuila | KY952307 |
| Nyctaginaceae | *Acleisanthes purpusiana*<br>(Heimerl) R.A.Levin | Billie L. Turner 6205 (TEX) | Mexico: Coahuila | KY952308 |
| Nyctaginaceae | *Acleisanthes undulata*<br>(B.A.Fowler & B.L.Turner) R.A.Levin | James Henrickson 23195 (TEX) | Mexico: Coahuila | KY952310 |
| Nyctaginaceae | *Acleisanthes wrightii*<br>(A.Gray) Benth. & Hook. | Michael J. Moore et al. 620 (OC) | United States: Texas | KY952311 |
| Nyctaginaceae | *Allionia choisyi* Standl. | Norman A. Douglas 2187 (DUKE) | Mexico: Coahuila | KY952315 |
| Nyctaginaceae | *Allionia incarnata* L. | Michael J. Moore et al. 1352 (OC) | Mexico: Nuevo Leon | KY952316 |
| Nyctaginaceae | *Allionia* sp. | Michael J. Moore 424 (OC) | United States: Texas | KY952317 |
| Nyctaginaceae | *Andradea floribunda*<br>Allemão | André M. Amorim 2294 (NY) | Brazil | KY952324 |
| Nyctaginaceae | *Andradea floribunda*<br>Allemão | Jacquelyn Ann Kallunki 701 (NY) | Brazil | KY952325 |
| Nyctaginaceae | *Anulocaulis annulatus* | Richard W. | United States: | KY952326 |
|  | (Coville) Standl. | Spellenberg 3162 (NMC) | California |  |
| Nyctaginaceae | *Anulocaulis eriosolenus* (A.Gray) Standl. | James Henrickson et al. 23103 (TEX) | Mexico: Coahuila | KY952327 |
| Nyctaginaceae | *Anulocaulis eriosolenus* (A.Gray) Standl. | Michael J. Moore et al. 611 (OC) | United States: Texas | KY952328 |
| Nyctaginaceae | *Anulocaulis hintoniorum* B.L.Turner | Patricia Hernández Ledesma 52 (MEXU) | Mexico: Coahuila | KY952329 |
| Nyctaginaceae | *Anulocaulis leiosolenus* (Torr.) Standl. var. *gyssogenus* (Waterf.) Spellb. & T.Wootten | Michael J. Moore 402 (OC) | United States: New Mexico | KY952330 |
| Nyctaginaceae | *Anulocaulis leiosolenus* (Torr.) Standl. var. *howardii* Spellb. & T.Wootten | Thomas Wootten et al. s.n. (NMC) | United States: New Mexico | KY952331 |
| Nyctaginaceae | *Anulocaulis leiosolenus* (Torr.) Standl. var. *lasianthus* I.M.Johnston | Michael J. Moore et al. 610 (OC) | United States: Texas | KY952332 |
| Nyctaginaceae | *Anulocaulis leiosolenus* (Torr.) Standl. var. *leiosolenus* | Michael J. Moore 493 (OC) | United States: Texas | KY952333 |
| Nyctaginaceae | *Anulocaulis leiosolenus* (Torr.) Standl. var. *leiosolenus* | Michael J. Moore et al. 825 (OC) | United States: Nevada | KY952334 |
| Nyctaginaceae | *Anulocaulis leiosolenus* (Torr.) Standl. var. *leiosolenus* | Michael J. Moore et al. 853 (OC) | United States: Arizona | KY952335 |
| Nyctaginaceae | *Anulocaulis reflexus* I.M.Johnst. | Michael J. Moore et al. 242 (TEX) | Mexico: Chihuahua | KY952336 |
| Nyctaginaceae | *Anulocaulis reflexus* I.M.Johnst. | Michael J. Moore 483 (OC) | United States: Texas | KY952337 |
| Nyctaginaceae | *Boerhavia anisophylla* Torr. | Norman A. Douglas 2194 (DUKE) | Mexico: Durango | KY952341 |
| Nyctaginaceae | *Boerhavia ciliata* Brandegees | Norman A. Douglas 2145 (DUKE) | United States: Texas | KY952342 |
| Nyctaginaceae | *Boerhavia coccinea* Mill. | Michael J. Moore 366 (OC) | United States: New Mexico | KY952343 |
| Nyctaginaceae | *Boerhavia coulteri* (Hook.f.) S.Watson var. *palmeri* (S.Watson) Spellb. | Richard W. Spellenberg 13273 (NMC) | United States: Arizona | KY952344 |
| Nyctaginaceae | *Boerhavia dominii* Meikle & Hewson | H. Smyth 42 (NY) | Australia: South Australia | KY952345 |
| Nyctaginaceae | *Boerhavia gracillima* Heimerl | Richard W. Spellenberg 12447 (NMC) | United States: Texas | KY952347 |
| Nyctaginaceae | *Boerhavia intermedia* M.E.Jones | Richard W. Spellenberg 13279 | United States: Arizona | KY952348 |
|  |  | (NMC) |  |  |
| Nyctaginaceae | *Boerhavia lateriflora* Standl. | Norman A. Douglas 2161 (DUKE) | Mexico: Sonora | KY952349 |
| Nyctaginaceae | *Boerhavia linearifolia* A.Gray | Michael J. Moore et al. 581 (OC) | United States: Texas | KY952350 |
| Nyctaginaceae | *Boerhavia purpurascens* A.Gray | Richard W. Spellenberg 13261 (NMC) | United States: Arizona | KY952351 |
| Nyctaginaceae | *Boerhavia repens* L. | J. S. Rose 2 | United States: Hawaii | KY952352 |
| Nyctaginaceae | *Boerhavia repens* L. | Richard W. Spellenberg 7183 (NMC) | Yemen: Sana | KY952353 |
| Nyctaginaceae | *Boerhavia* sp. | Erin Tripp et al. 4090 (OC) | Namibia | KY952346 |
| Nyctaginaceae | *Boerhavia torreyana* (S.Watson) Standl. | Michael J. Moore et al. 633 (OC) | United States: Texas | KY952354 |
| Nyctaginaceae | *Bougainvillea campanulata* Heimerl | Michael Nee 51257 (TEX) | Bolivia: Santa Cruz | KY952355 |
| Nyctaginaceae | *Bougainvillea glabra* Choisy | Michael J. Moore 538 (OC) | United States: Ohio (cultivated) | KY952356 |
| Nyctaginaceae | *Bougainvillea spinosa* (Cav.) Heimerl | J. Saunders et al. 3371 (TEX) | Argentina: San Juan | KY952357 |
| Nyctaginaceae | *Bougainvillea stipitata* Griseb. | Michael Nee 50723 (TEX) | Bolivia: Santa Cruz | KY952358 |
| Nyctaginaceae | *Colignonia glomerata* Griseb. | Michael Nee 52523 (NY) | Bolivia | KY952364 |
| Nyctaginaceae | *Colignonia scandens* Benth. | Martin Grantham 63 (SFBG living collection, accession 1996-0202) | Ecuador | KY952365 |
| Nyctaginaceae | *Commicarpus ambiguus* Meikle | Mats Thulin 11015 (UPS) | Somalia: Sanaag | KY952366 |
| Nyctaginaceae | *Commicarpus arabicus* Meikle | Mats Thulin et al. 9294 (UPS) | Yemen: Taizz | KY952367 |
| Nyctaginaceae | *Commicarpus arabicus* Meikle | Richard W. Spellenberg 7217 (NMC) | Yemen: Ibb | KY952368 |
| Nyctaginaceae | *Commicarpus arabicus* Meikle | Richard W. Spellenberg 7297 (NMC) | Yemen: Ibb | KY952369 |
| Nyctaginaceae | *Commicarpus australis* (Meikle) Govaerts | Richard W. Spellenberg et al. 9469 (NMC) | Australia: Western Australia | KY952370 |
| Nyctaginaceae | *Commicarpus boissieri* (Heimerl) Cufod. | Mats Thulin 11423 (UPS) | Oman: Dhofar | KY952371 |
| Nyctaginaceae | *Commicarpus boissieri* (Heimerl) Cufod. | Carl J. Rothfels et al. 4331 | Oman: Ash Sharqiyah | KY952373 |
| Nyctaginaceae | *Commicarpus* | Patricia Hernández | Mexico: Baja | KY952372 |
|  | *brandegeei* Standl. | Ledesma 55 (MEXU) | California Sur |  |
| Nyctaginaceae | *Commicarpus coctoris*<br>N.A.Harriman | Richard W.<br>Spellenberg et al.<br>12883 (NMC) | Mexico:<br>Oaxaca | KY952374 |
| Nyctaginaceae | *Commicarpus commersonii* (Baill.)<br>Cavaco | Mats Thulin et al.<br>11836 (UPS) | Madagascar:<br>Toliara | KY952380 |
| Nyctaginaceae | *Commicarpus decipiens*<br>Meikle | Erin Tripp et al. 4127<br>(NMC) | Namibia | KY952375 |
| Nyctaginaceae | *Commicarpus grandiflorus* (A.Rich.)<br>Standl. | Mats Thulin et al.<br>9311 (UPS) | Yemen: Taizz | KY952376 |
| Nyctaginaceae | *Commicarpus greenwayi*<br>Meikle | Mats Thulin 606<br>(UPS) | Tanzania:<br>Iringa | KY952377 |
| Nyctaginaceae | *Commicarpus helenae*<br>(Roem. & Schult.)<br>Meikle | Richard W.<br>Spellenberg et al.<br>7504 (NMC) | Yemen:<br>Dhamar | KY952378 |
| Nyctaginaceae | *Commicarpus hiranensis*<br>Thulin | Mats Thulin et al.<br>11225 (UPS) | Ethiopia:<br>Harerge | KY952379 |
| Nyctaginaceae | *Commicarpus mistus*<br>Thulin | Mats Thulin et al.<br>9786 (UPS) | Yemen:<br>Mahrah | KY952381 |
| Nyctaginaceae | *Commicarpus parviflorus* Thulin | Mats Thulin 6318<br>(UPS) | Somalia:<br>Banaadir | KY952382 |
| Nyctaginaceae | *Commicarpus pedunculatus* (A.Rich.)<br>Cufod. | Mats Thulin 1301<br>(UPS) | Ethiopia:<br>Arussi | KY952383 |
| Nyctaginaceae | *Commicarpus plumbagineus* (Cav.)<br>Standl. | Mats Thulin 10747<br>(UPS) | Somalia:<br>Togdheer | KY952384 |
| Nyctaginaceae | *Commicarpus plumbagineus* (Cav.)<br>Standl. | Mats Thulin et al.<br>11330 (UPS) | Ethiopia:<br>Harerge | KY952385 |
| Nyctaginaceae | *Commicarpus plumbagineus* (Cav.)<br>Standl. | Richard W.<br>Spellenberg et al.<br>7374 (NMC) | Yemen: Ta'izz | KY952386 |
| Nyctaginaceae | *Commicarpus praetermissus*<br>N.A.Harriman | Richard W.<br>Spellenberg et al.<br>12905 (NMC) | Mexico:<br>Michoacán | KY952387 |
| Nyctaginaceae | *Commicarpus reniformis*<br>(Chiov.) Cufod. | Mats Thulin 4200<br>(UPS) | Somalia: Sool | KY952388 |
| Nyctaginaceae | *Commicarpus reniformis*<br>(Chiov.) Cufod. | Mats Thulin et al.<br>8337 (UPS) | Yemen:<br>Hadramaut | KY952389 |
| Nyctaginaceae | *Commicarpus scandens*<br>(L.) Standl. | Michael J. Moore<br>1127 (OC) | United States:<br>Texas | KY952390 |
| Nyctaginaceae | *Commicarpus scandens*<br>(L.) Standl. | Richard W.<br>Spellenberg et al.<br>12887 (NMC) | Mexico:<br>Puebla | KY952391 |
| Nyctaginaceae | *Commicarpus sinuatus*<br>Meikle | Mats Thulin 10737<br>(UPS) | Somalia:<br>Woqooyi<br>Galbeed | KY952392 |
| Nyctaginaceae | *Commicarpus sinuatus*<br>Meikle | Richard W.<br>Spellenberg 7144<br>(NMC) | Yemen:<br>Sana'a | KY952393 |
| Nyctaginaceae | *Commicarpus sinuatus*<br>Meikle | Richard W.<br>Spellenberg 7506<br>(NMC) | Yemen:<br>Dhamar | KY952394 |
| Nyctaginaceae | *Commicarpus squarrosus* (Heimerl)<br>Standl. var. *squarrosus* | Erin Tripp et al. 4049<br>(NMC) | Namibia | KY952395 |
| Nyctaginaceae | *Commicarpus stenocarpus* (Chiov.)<br>Cufod. | Mats Thulin et al.<br>8062 (UPS) | Yemen:<br>Hadramaut | KY952396 |
| Nyctaginaceae | *Cuscatlania vulcanicola*<br>Standl. | José L. Linares 12938<br>(MEXU) | El Salvador:<br>Sonsonate | KY952397 |
| Nyctaginaceae | *Cuscatlania vulcanicola*<br>Standl. | José L. Linares 13440<br>(MEXU) | El Salvador:<br>Sonsonate | KY952398 |
| Nyctaginaceae | *Cyphomeris gypsophiloides*<br>(M.Martens & Galeotti)<br>Standl. | Michael J. Moore et<br>al. 582 (OC) | United States:<br>Texas | KY952399 |
| Nyctaginaceae | *Grajalesia fasciculata*<br>(Standl.) Miranda | José L. Linares 13416<br>(MEXU) | El Salvador:<br>Sonsonate | KY952409 |
| Nyctaginaceae | *Guapira discolor*<br>(Spreng.) Little | Richard W.<br>Spellenberg 13294<br>(NMC) | United States:<br>Florida | KY952410 |
| Nyctaginaceae | *Guapira eggersiana*<br>(Heimerl) Lundell | Scott A. Mori<br>25542/40 (NY) | French Guiana | KY952411 |
| Nyctaginaceae | *Mirabilis albida*<br>(Walter) Heimerl | Norman A. Douglas<br>2035 (DUKE) | United States:<br>Arizona | KY952416 |
| Nyctaginaceae | *Mirabilis albida*<br>(Walter) Heimerl | William R. Carr<br>11075 (TEX) | United States:<br>Texas | KY952417 |
| Nyctaginaceae | *Mirabilis alipes*<br>(S.Watson) Pilz | Arnold Tiehm 13461<br>(TEX) | United States:<br>Nevada | KY952418 |
| Nyctaginaceae | *Mirabilis bigelovii*<br>A.Gray var. *retrorsa* (A.<br>Heller) Munz | James D. Morefield<br>et al. 3780 (TEX) | United States:<br>California | KY952419 |
| Nyctaginaceae | *Mirabilis* cf. *glabrifolia*<br>(Ortega) I.M.Johnst. | Michael J. Moore et<br>al. 1244 (OC) | Mexico: San<br>Luis Potosi | KY952428 |
| Nyctaginaceae | *Mirabilis* cf. *nesomii*<br>B.L.Turner | George S. Hinton<br>25567 (TEX) | Mexico:<br>Nuevo Leon | KY952449 |
| Nyctaginaceae | *Mirabilis coccinea*<br>(Torr.) Benth. & Hook.f. | Norman A. Douglas<br>2133 (DUKE) | United States:<br>Arizona | KY952420 |
| Nyctaginaceae | *Mirabilis coccinea*<br>(Torr.) Benth. & Hook.f. | Steven P.<br>McLaughlin et al.<br>9354 (ARIZ) | United States:<br>Arizona | KY952421 |
| Nyctaginaceae | *Mirabilis comata*<br>(Small) Standl. | Norman A. Douglas<br>2084 (DUKE) | United States:<br>Arizona | KY952422 |
| Nyctaginaceae | *Mirabilis decumbens*<br>(Nutt.) Daniels | Richard W.<br>Spellenberg et al.<br>4073 (TEX) | Mexico:<br>Zacatecas | KY952423 |
| Nyctaginaceae | *Mirabilis donahooiana*<br>Le Duc | Alice Le Duc et al.<br>247 (TEX) | Mexico:<br>Michoacán | KY952424 |
| Nyctaginaceae | *Mirabilis exserta*<br>Brandege | Pedro Tenorio 10586<br>(MEXU) | Mexico | KY952425 |
| Nyctaginaceae | *Mirabilis gigantea*<br>(Standl.) Shinnars | J. Quayle et al. 752<br>(TEX) | United States:<br>Texas | KY952426 |
| Nyctaginaceae | *Mirabilis glabra*<br>(S.Watson) Standl. | Michael J. Moore et<br>al. 674 (OC) | United States:<br>New Mexico | KY952446 |
| Nyctaginaceae | *Mirabilis glabrifolia*<br>(Ortega) I.M.Johnst. | Guy Nesom et al.<br>7654 (TEX) | Mexico:<br>Coahuila | KY952427 |
| Nyctaginaceae | *Mirabilis glabrifolia*<br>(Ortega) I.M.Johnst. | Michael J. Moore et<br>al. 1325 (OC) | Mexico:<br>Nuevo Leon | KY952429 |
| Nyctaginaceae | *Mirabilis gracilis*<br>(Standl.) LeDuc | Alice Le Duc et al. 71<br>(TEX) | Mexico:<br>Jalisco | KY952430 |
| Nyctaginaceae | *Mirabilis grandiflora*<br>(Standl.) Standl. | EDL 1863 (MEXU) | Mexico | KY952431 |
| Nyctaginaceae | *Mirabilis greenii*<br>S.Watson | George E. Pilz 998<br>(TEX) | United States:<br>California | KY952432 |
| Nyctaginaceae | *Mirabilis himalaica*<br>(Edgew.) Heimerl var.<br>*chinensis* Heimerl | D. E. Boufford et al.<br>32449 (F ) | China: Xizang<br>(Tibet) | KY952433 |
| Nyctaginaceae | *Mirabilis himalaica*<br>(Edgew.) Heimerl var.<br>*chinensis* Heimerl | D. E. Boufford et al.<br>41198 (F) | China: Xizang<br>(Tibet) | KY952434 |
| Nyctaginaceae | *Mirabilis himalaica*<br>(Edgew.) Heimerl var.<br>*chinensis* Heimerl | D. E. Boufford et al.<br>41435 (F ) | China: Xizang<br>(Tibet) | KY952435 |
| Nyctaginaceae | *Mirabilis hintoniorum*<br>Le Duc | Patricia Hernández<br>Ledesma 118<br>(MEXU) | Mexico:<br>Michoacán | KY952436 |
| Nyctaginaceae | *Mirabilis jalapa* L. | Michael J. Moore s.n. | United States<br>(cultivated) | KY952437 |
| Nyctaginaceae | *Mirabilis laevis* (Benth.)<br>Curran | Andrew C. Sanders et<br>al. 29410 (TEX) | United States:<br>California | KY952438 |
| Nyctaginaceae | *Mirabilis latifolia*<br>(A.Gray) Diggs,<br>Lipscomb & O'Kennon | Victor L. Cory 24549<br>(GH) | United States:<br>Texas | KY952439 |
| Nyctaginaceae | *Mirabilis linearis*<br>(Pursh) Heimerl | Billie L. Turner 21-854 (TEX) | United States:<br>Texas | KY952440 |
| Nyctaginaceae | *Mirabilis linearis*<br>(Pursh) Heimerl var.<br>*decipiens* (Standl.)<br>S.L.Welsh | Michael J. Moore et<br>al. 1984 (OC) | Mexico:<br>Coahuila | KY952441 |
| Nyctaginaceae | *Mirabilis longiflora* L. | Michael J. Moore et<br>al. 1230 (OC) | Mexico: San<br>Luis Potosi | KY952442 |
| Nyctaginaceae | *Mirabilis longiflora* L.<br>var. *wrightiana* (A.Gray<br>ex Britton & Kearney)<br>Kearney & Peebles | Alice Le Duc 185<br>(TEX) | United States:<br>New Mexico | KY952443 |
| Nyctaginaceae | *Mirabilis melanotricha* | Michael J. Moore et | Mexico: San | KY952444 |
|  | (Standl.) Spellenb. | al. 1191 (OC) | Luis Potosi |  |
| Nyctaginaceae | *Mirabilis melanotricha* (Standl.) Spellenb. | Norman A. Douglas 2067 (DUKE) | United States: New Mexico | KY952445 |
| Nyctaginaceae | *Mirabilis multiflora* (Torr.) A.Gray | Michael J. Moore 1110 (OC) | United States: Texas | KY952447 |
| Nyctaginaceae | *Mirabilis multiflora* (Torr.) A.Gray | Norman A. Douglas 2037 (DUKE) | United States: Arizona | KY952448 |
| Nyctaginaceae | *Mirabilis nesomii* B.L.Turner | Michael J. Moore et al. 2179 (NMC) | Mexico: Nuevo Leon | KY952450 |
| Nyctaginaceae | *Mirabilis nesomii* B.L.Turner | Michael J. Moore et al. 2643 (NMC) | Mexico: Nuevo Leon | KY952451 |
| Nyctaginaceae | *Mirabilis nyctaginea* (Michx.) MacMill. | William R. Carr 14590 (TEX) | United States: Texas | KY952452 |
| Nyctaginaceae | *Mirabilis oligantha* (Standl.) Standl. | José L. Panero 2816 (MEXU) | Mexico: Baja California | KY952453 |
| Nyctaginaceae | *Mirabilis oxybaphoides* (A.Gray) A.Gray | George S. Hinton 25572 (TEX) | Mexico: Nuevo Leon | KY952454 |
| Nyctaginaceae | *Mirabilis polonii* Le Duc | Alice Le Duc 259 (MEXU) | Mexico: Nuevo Leon | KY952455 |
| Nyctaginaceae | *Mirabilis pringlei* Weath. | Alice Le Duc et al. 63 (TEX) | Mexico: Jalisco | KY952456 |
| Nyctaginaceae | *Mirabilis pudica* Barneby | Arnold Tiehm 10971 (TEX) | United States: Nevada | KY952457 |
| Nyctaginaceae | *Mirabilis texensis* (J.M.Coult.) B.L.Turner | Billie L. Turner 22-417 (TEX) | United States: Texas | KY952458 |
| Nyctaginaceae | *Mirabilis triflora* Benth. | Ramón Cuevas G. et al. 3415 (MEXU) | Mexico: Jalisco | KY952459 |
| Nyctaginaceae | *Mirabilis urbani* Heimerl | Mark Fishbein et al. 5107 (MEXU) | Mexico: Michoacan | KY952460 |
| Nyctaginaceae | *Mirabilis violacea* (L.) Heimerl | Patricia Hernández Ledesma 63 (MEXU) | Mexico: Distrito Federal | KY952461 |
| Nyctaginaceae | *Mirabilis viscosa* Cav. | Michael J. Moore et al. 1824 (NMC) | Mexico: San Luis Potosi | KY952462 |
| Nyctaginaceae | *Mirabilis viscosa* Cav. | Patricia Hernández Ledesma 13 (MEXU) | Mexico | KY952463 |
| Nyctaginaceae | *Neea belizensis* Donn.Sm. | Cyrus L. Lundell 17692 (TEX) | Guatemala: Petén | KY952465 |
| Nyctaginaceae | *Neea cauliflora* Poepp. & Endl. | Schanke S15106 (NY) | Peru | KY952466 |
| Nyctaginaceae | *Neea psychotrioides* Donn.Sm. | Robert L. Wilbur 63654 | Costa Rica: Heredia | KY952467 |
| Nyctaginaceae | *Nyctaginia capitata* Choisy | Michael J. Moore et al. 617 (OC) | United States: Texas | KY952470 |
| Nyctaginaceae | *Okenia hypogaea* Schltdl. & Cham. | Thomas R. Van Devender et al. 92-1069 (NMC) | Mexico: Sonora | KY952471 |
| Nyctaginaceae | *Pisonia aculeata* L. | C. Martínez 1209 (TEX) | Mexico: Oaxaca | KY952483 |
| Nyctaginaceae | *Pisonia brunoniana* | J. S. Rose 3 | United States: | KY952484 |
|  | Endl. |  | Hawaii |  |
| Nyctaginaceae | *Pisonia capitata*<br>(S.Watson) Standl. | Ana L. Reina<br>Guerrero et al. 2000-193 (NMC) | Mexico:<br>Sonora | KY952485 |
| Nyctaginaceae | *Pisonia capitata*<br>(S.Watson) Standl. | Thomas R. Van<br>Devender et al. 2003-17 (TEX) | United States:<br>Arizona | KY952486 |
| Nyctaginaceae | *Pisonia macranthocarpa*<br>(Donn.Sm.) Donn.Sm. | Dennis E. Breedlove<br>et al. 30361 (TEX) | Mexico:<br>Chiapas | KY952487 |
| Nyctaginaceae | *Pisonia sandwicensis*<br>Hillebr. | Flora K. Samis 1<br>(Lyon Arboretum<br>living collection) | United States:<br>Hawaii | KY952488 |
| Nyctaginaceae | *Pisonia sylvatica* Standl. | José L. Linares 13403<br>(MEXU) | El Salvador:<br>Sonsonate | KY952489 |
| Nyctaginaceae | *Pisonia umbellifera*<br>(J.R.Forst. & G.Forst.)<br>Seem. | Flora K. Samis 12<br>(Lyon Arboretum<br>living collection,<br>accession 68.0453) | United States:<br>Hawaii | KY952490 |
| Nyctaginaceae | *Pisonia zapallo* Griseb. | Israel G. Vargas et al.<br>2001 (TEX) | Bolivia: Santa<br>Cruz | KY952491 |
| Nyctaginaceae | *Pisoniella arborescens*<br>(Lag. & Rodr.) Standl. | Alice Le Duc et al.<br>231 (NMC) | Mexico:<br>Oaxaca | KY952492 |
| Nyctaginaceae | *Pisoniella arborescens*<br>(Lag. & Rodr.) Standl. | William R. Anderson<br>13522 (NY) | Mexico:<br>Oaxaca | KY952493 |
| Nyctaginaceae | *Ramisia brasiliensis*<br>Oliv. | Jomar G. Jardim<br>1507 (NY) | Brazil | KY952495 |
| Nyctaginaceae | *Reichenbachia hirsuta*<br>Spreng. | Michael Nee 47813<br>(NY) | Bolivia | KY952496 |
| Nyctaginaceae | *Reichenbachia*<br>*paraguayensis*<br>(D.Parodi) Dugand &<br>Daniel | Maria Maguidaura<br>Hatschbach 49218<br>(NY) | Brazil | KY952497 |
| Nyctaginaceae | *Salpianthus arenarius*<br>Bonpl. | Richard W.<br>Spellenberg 12903<br>(NMC) | Mexico:<br>Michoacán | KY952503 |
| Nyctaginaceae | *Salpianthus*<br>*macrodonatus* Standl. | Thomas R. Van<br>Devender et al. 91-894 (NMC) | Mexico:<br>Sonora | KY952504 |
| Nyctaginaceae | *Salpianthus*<br>*purpurascens* (Cav. ex<br>Lag.) Hook. & Arn. | Richard W.<br>Spellenberg et al.<br>12885 (NMC) | Mexico:<br>Oaxaca | KY952505 |
| Nyctaginaceae | *Tripterocalyx carneus*<br>(Greene) L.A.Galloway | Norman A. Douglas<br>2060 (DUKE) | United States:<br>New Mexico | KY952525 |
| Nyctaginaceae | *Tripterocalyx crux-*<br>*maltae* (Kellogg) Standl. | Arnold Tiehm et al.<br>12213 (TEX) | United States:<br>Nevada | KY952526 |
| Nyctaginaceae | *Tripterocalyx*<br>*micranthus* (Torr.)<br>Hook. | B. MacLeod et al.<br>751 (TEX) | United States:<br>Colorado | KY952527 |
| Phytolaccaceae | *Agdestis clematidea*<br>Moc. & Sessé ex DC. | George S. Hinton<br>25023 (TEX) | Mexico:<br>Tamaulipas | KY952313 |
| Phytolaccaceae | *Gallesia integrifolia* (Spreng.) Harms | Michael Nee et al. 50072 (TEX) | Bolivia: Santa Cruz | KY952407 |
| Phytolaccaceae | *Hillieria latifolia* (Lam.) H.Walter | Michael Nee 33807 (TEX) | Bolivia: Santa Cruz | KY952413 |
| Phytolaccaceae | *Petiveria alliacea* L. | Lucas C. Majure 4132 (FLAS) | United States: Florida | KY952476 |
| Phytolaccaceae | *Phytolacca americana* L. | Michael J. Moore 342 (OC) | United States: Ohio | KY952480 |
| Phytolaccaceae | *Phytolacca icosandra* L. | Mark H. Mayfield et al. 1001 (TEX) | Mexico: Guerrero | KY952481 |
| Phytolaccaceae | *Phytolacca octandra* L. | Juan A. Encina et al. 1545 (TEX) | Mexico: Nuevo Leon | KY952482 |
| Phytolaccaceae | *Rivina humilis* L. | Michael J. Moore 1129 (OC) | United States: Texas | KY952499 |
| Phytolaccaceae | *Seguieria aculeata* Jacq. | Elsa Zardini et al. 22101 (TEX) | Paraguay | KY952510 |
| Phytolaccaceae | *Seguieria paraguariensis* Morong | Michael Nee 48735 (TEX) | Bolivia: Santa Cruz | KY952511 |
| Phytolaccaceae | *Trichostigma octandrum* (L.) H.Walter | Michael Nee 47094 (TEX) | Bolivia: Santa Cruz | KY952522 |
| Phytolaccaceae | *Trichostigma peruvianum* (Moq.) H.Walter | Flora K. Samis 10 (Lyon Arboretum living collection, accession 94.0377) | United States: Hawaii | KY952523 |
| Plumbaginaceae | *Aegialitis annulata* R.Br. | Christopher T. Martine 4043 (OC) | Australia: Western Australia | KY952312 |
| Plumbaginaceae | *Limonium limbatum* Small | Michael J. Moore et al. 694 (OC) | United States: New Mexico | KY952414 |
| Plumbaginaceae | *Plumbago scandens* L. | Michael J. Moore et al. 1828 (OC) | Mexico: San Luis Potosi | KY952494 |
| Polygonaceae | *Eriogonum longifolium* Nutt. var. *longifolium* | Michael J. Moore 1796 (OC) | United States: Texas | KY952404 |
| Polygonaceae | *Eriogonum rotundifolium* Benth. | Michael J. Moore 1769 (OC) | United States: New Mexico | KY952405 |
| Polygonaceae | *Persicaria odorata* LaLlave | Flora K. Samis 9 (Lyon Arboretum living collection, accession 88.0439) | United States: Hawaii | KY952475 |
| Polygonaceae | *Persicaria* sp. | Michael J. Moore 1177 | United States: Ohio | KY952474 |
| Polygonaceae | *Reynoutria japonica* (Houtt.) Ronse Decr. | Michael J. Moore 2188 (OC) | United States: Ohio | KY952498 |
| Polygonaceae | *Rumex albens* Hillebr. | Flora K. Samis 4 (Lyon Arboretum living collection, accession 2008-0119) | United States: Hawaii | KY952500 |
| Polygonaceae | *Rumex* sp. | Michael J. Moore 1800 (OC) | United States: Texas | KY952501 |
| Polygonaceae | *Rumex* sp. | Michael J. Moore | United States: | KY952502 |
|  |  | 1805 (OC) | Texas |  |
| Sarcobataceae | *Sarcobatus vermiculatus* (Hook.) Torr. | Michael J. Moore et al. 813 (OC) | United States: Utah | KY952508 |
| Stegnospermataceae | *Stegnosperma cubense* A.Rich. | Silvia H. Salas Morales 2649 (NY) | Mexico: Oaxaca | KY952513 |
| Talinaceae | *Talinum* cf. *aurantiacum* Engelm. | Michael J. Moore et al. 1985 (OC) | Mexico: Coahuila | KY952517 |
| Talinaceae | *Talinum fruticosum* (L.) Juss. | Flora K. Samis 8 (Lyon Arboretum living collection, accession 2012.0008) | United States: Hawaii | KY952518 |
| Talinaceae | *Talinum paniculatum* (Jacq.) Gaertn. | Michael J. Moore 1789 (OC) | United States (cultivated) | KY952520 |
| Talinaceae | *Talinum* sp. | Michael J. Moore et al. 1974 (MEXU) | Mexico: Coahuila | KY952519 |

**Table 2:**
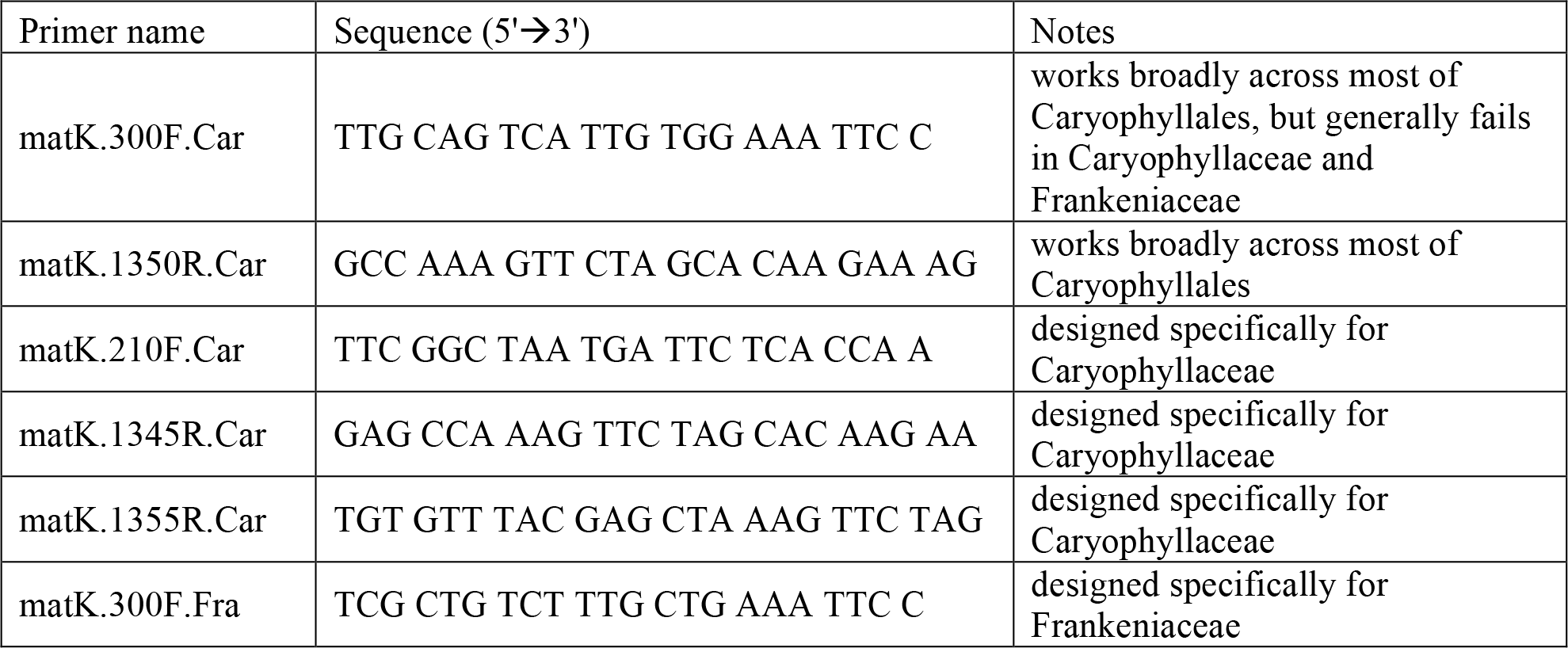
List of primers used to amplify the *matK* sequences newly reported here. Within each primer name, the number indicates the approximate position of the primer in nucleotides downstream from the start of *matK*.

### Molecular Data for Phylogenetic Reconstruction

Nucleotide data from the nuclear ribosomal internal transcribed spacers (ITS) and *phyC* gene, and the plastid loci *matK*, *ndhF, rbcL*, *trnH-psbA* spacer, and *trnL-trnF* spacer were used to reconstruct the phylogeny. These data were gathered first using PHLAWD (Smith & Donoghue, 2008; Smith *et al*., 2009) and then curated and combined with newly sequenced *matK* data for 124 additional species. This yielded the following sampling: ITS 2,969 species, *matK* 2,270 species, *ndhF* 417 species, *phyC* 172 species, *rbcL* 947 species, *trnH-psbA* 240 species, and *trnL-trnF* 1,996 species. We used *mat*K, *rbc*L, and *ndh*F sequences from *Aextoxicon, Apium, Berberidopsis, Campanula, Clethra, Coffea, Echinops, Helwingia, Ilex, Ipomoea, Lamium, Lonicera, Nyssa, Polysoma, Primula, Santalum, Valeriana*, and *Viburnum* to represent outgroups.

### Phylogenetic Reconstruction

We conducted phylogenetic analyses with RAxML v7.2.8 (Stamatakis, 2014) using the full analysis command, -f a, which conducts a rapid bootstrap and then a full maximum likelihood search. The combined bootstrap and maximum likelihood search allows for a more thorough maximum likelihood analysis where the initial rapid bootstrap results prime the maximum likelihood analysis. However, we did not use the rapid bootstrap trees from this analysis and instead, we conducted a full bootstrap, generating the bootstrap dataset using phyx (Brown *et al*., 2017) and then conducting individual maximum likelihood runs on each constructed bootstrap dataset. This allowed us to conduct SH-like approximate likelihood ratio test (SH-aLRT; Guindon *et al*., 2010) on the resulting bootstrap set. We conducted bootstraps within gene regions and we retained the individual bootstrap alignments to conduct additional analyses (i.e., bootstrapped alignments contained the same number of gene-specific sites as the empirical alignment). On each of the resulting trees of the bootstrap and the maximum likelihood tree, we conducted SH-aLRTs as implemented in RAxML. These analyses calculate support for each edge while also finding the NNI-optimal topology. RAxML completed the likelihood search for each of these bootstrap replicates, however the SH-aLRT analyses often resulted in an improved maximum likelihood topology. The trees that resulted from the SH-aLRT, ML, and bootstrap samples, were used for further analyses. Because several deep relationships within Caryophyllales are hard to resolve without large amounts of molecular data that are unavailable for most of the taxa included in this analysis (Yang *et al*., 2015), for all phylogenetic analyses we applied the following topological constraint: (Droseraceae, (*Microtea*, (Stegnospermataceae, Limeaceae, (Lophiocarpaceae, (Barbeuiaceae, Aizoaceae))))) as per previous analysis (Brockington *et al*., 2009; Yang *et al*., 2015).

### Divergence Time Estimation

Few tractable options for divergence time estimation exist for datasets of the size presented here. We use the penalized likelihood approach (Sanderson, 2003) as implemented in the program treePL (Smith & O’Meara, 2012), which can handle large-scale phylogenies. The early fossil record of the Caryophyllales is sparse with only a few known records (Friis *et al*., 2011; Arakaki *et al*., 2011): (1) fossil pollen has been ascribed to Amaranthaceae (*Chenopodipollis*) from the Paleocene of Texas (Nichols & Traverse, 1971); (2) a putative fossil infructescence from within the Phytolaccaceae in the Campanian has also been reported (Cevallos-Ferriz *et al*., 2008), but this phylogenetic position has been disputed (pers. comm. S. Manchester) and hence we excluded it; (3) Jordan & Macphail (2003) describe a middle to late Eocene inflorescence from the species *Caryophylloflora paleogenica*, ascribed to Caryophyllaceae; (4) pollen from Argentina within the Nyctaginaceae has been reported from the middle Eocene (Zetter *et al*., 1999); and (5) fossil pollen and seeds of *Aldrovanda* (Degreef, 1997). The penalized likelihood method performs better when a calibration is used at the root. For this calibration, and because there is no fossil record for the earliest Caryophyllales, we use a secondary calibration from the comprehensive angiosperm divergence time analyses of Bell *et al*. (2010). We attached several other secondary calibrations to major clades where fossils are not available (Ocampo & Columbus 2010; Arakaki *et al*., 2011; Schuster *et al*., 2013; Valente *et al*., 2013; see Supp. Table S1 for detail on placement and calibrations). We conducted a priming analysis to determine the best optimization parameter values. We then performed a cross validation analysis using the random cross validation setting to determine the optimal smoothing parameter value.

### Climate occupancy analyses

We downloaded 6,592,700 georeferenced occurrences for the Caryophyllales from GBIF (accessed on 6/1/2015; http://gbif.org). After removing samples present in living collections, and therefore not necessarily representative of native climates, and removing samples whose localities were over water, 6,009,552 samples remained. We extracted bioclimatic values for each coordinate using the 2.5 arc-minute resolution data from WorldClim (http://worldclim.org). We only included taxa that had at least three samples in these analyses to reduce potential errors and to have the minimum number of samples required to calculate mean and variance. The resulting overlap of the taxa represented in both the geographic and genetic data was 2,843 taxa. We conducted principal component analyses (PCA) on these extracted values. With both the bioclimatic values and the first two axes of the PCA, we conducted ancestral state reconstruction analyses.

We also conducted contrast analyses and calculated Brownian motion rates of evolution between sister clades (comparing duplicated lineages with their sisters) for mean annual precipitation, mean annual temperature, and principal component axis 1. We calculated contrasts using phylogenetic independent contrasts. We calculated Brownian motion rates on sister lineages independently using the analytical solution for rate: 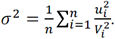

### Diversification analyses

To map diversification rate shifts, we conducted MEDUSA (Alfaro *et al*., 2009; Pennell *et al*., 2014) analyses on the maximum likelihood tree and the bootstrap trees. MEDUSA is far more computationally tractable than some other diversification estimation methods. Furthermore, we required the ability to feasibly integrate over the phylogenetic uncertainty within the phylogenetic dataset because of both the nature of the larger phylogenetic dataset and the inherent biological uncertainty within the Caryophyllales. MEDUSA fits a birth-death model of diversification (with parameters *r*: net diversification (birth - death), and ε: relative extinction (death / birth)) before using stepwise AIC (Burnham & Anderson, 2002) to identify shifts in rates of diversification. These complementary analyses accommodate topological and branch length uncertainty. We employed a birth-death model for 97 chronograms generated from nonparametric bootstrapping of the original matrix, inferring ML trees in RAxML, and estimating divergence times in treePL using the temporal constraints described above. We discarded three trees based on poor fossil placement resulting from phylogenetic uncertainty causing fossil placements to conflict.

### Whole Genome Duplication Identification

To identify WGDs (procedure described below), we generated a tree based on transcriptomic data. For this tree, we used 178 ingroup data sets (175 transcriptomes, 3 genomes) representing 169 species in 27 families and 40 outgroup genomes (Table S1-S2 in Yang et al. submitted). We mapped putative WGD events using multiple strategies: gene tree topology, plotting synonymous distance, and chromosome counts (Yang *et al*. 2015; Yang *et al*., submitted). For gene tree topology analyses, we performed two alternative strategies for mapping duplication events from gene trees to the species tree: mapping to the most recent common ancestor (MRCA), or mapping to the species tree only when gene tree and species tree topologies are compatible.

To conduct synonymous distance analyses, we performed the following procedure. For all ingroup Caryophyllales transcriptome data sets, we calculated the distribution of paralog synonymous distance following the same procedure as (Yang *et al*., 2015). We reduced highly similar peptide sequences with CD-HIT (-c 0.99 -n 5)(Li & Godzik 2006). We also carried out an all-by-all BLASTP within each taxon using an E value cutoff of 10 and -max_target_seq set to 20. Resulting hits with pident < 20% or niden < 50 amino acids were removed. We removed sequences with ten or more hits to avoid overrepresentation of gene families that experienced multiple recent duplications. We used the remaining paralog pairs and their corresponding CDS to calculate Ks values using the pipeline https://github.com/tanghaibao/bio-pipeline/tree/master/synonymous_calculation (accessed November 29, 2014). The pipeline first carries out pairwise protein alignment using default parameters in ClustalW (Larkin *et al*., 2007), back-translates the alignment to a codon alignment using PAL2NAL (Suyama *et al*., 2006), and calculates the synonymous substitution rate (Ks) using yn00 as part of the PAML package (Yang, 2007), with Nei–Gojobori correction for multiple substitutions (Nei & Gojobori, 1986). We obtained chromosome counts from the Chromosome Counts Database (CCDB; http://ccdb.tau.ac.il accessed Oct 5, 2015). When multiple counts were reported from different authors or different plants, we erred on the conservative estimate and recorded the lowest number. For species that were not available in the database, we found counts from the literature (e.g., Jepson eFlora http://ucjeps.berkeley.edu/eflora/ and Flora of North America http://floranorthamerica.org) or by a consensus from species of the same genera.

## Results and Discussion

### Phylogenetic results

Phylogenetic analyses showed strong support based on bootstrap and SH-aLRT values for the monophyly of most Caryophyllales families (see Fig. S1). We found strong support for the carnivorous clade including Droseraceae, Ancistrocladaceae, Nepenthaceae, Drosophyllaceae, and Dioncophyllaceae. There was also strong support for this clade as sister to a clade including Frankeniaceae, Tamaricaceae, Plumbaginaceae, and Polygonaceae. However, relationships among the families showed more varied support. There was weak support for the placement of other families relative to other early diverging Caryophyllales (see Fig. S1). There was strong support for Caryophyllaceae sister to Amaranthaceae. There was very weak support for Aizoaceae sister to Phytolaccaceae+Nyctaginaceae. As with previously published analyses, there was no support for the monophyly of Phytolaccaceae in the traditional sense (i.e., including Phytolaccaceae s.s., Petiveriaceae, and *Agdestis*; APG IV) and very weak support for the placement of Sarcobataceae. There was also weak support for the relationships among Limeaceae, Molluginaceae, and the Portulacineae. Many of these relationships have been found to be strongly supported but conflicting in different analyses (Brockington *et al*., 2009; Soltis *et al*., 2011; Yang *et al*., 2015; Smith *et al*., 2015; Walker *et al*., 2017). Here, we focused less on the systematic resolution within the Caryophyllales and instead examine the potential relationship of diversification and climate occupancy shifts to WGDs. Therefore, we placed more emphasis on including more taxa over that of more gene regions (i.e., transcriptomes) at the cost of more missing data. Confident resolution of many of the systematic relationships will require genomic and transcriptomic sampling, as well as more thorough taxon sampling (Yang *et al*., submitted).

### Climate occupancy reconstruction results

We performed climate occupancy ancestral reconstruction analyses on the phylogeny of 2,843 taxa that included taxa with at least three sampled geographic coordinates (Figs. 1-3). We conducted these analyses for visualization and for comparison with diversification and WGD results (see below). Results for individual bioclimatic variables and principal components can be found in Figs. S2-S4. Bioclimatic variable 1 (mean annual temperature, Fig. 1) showed that there are several strong phylogenetic patterns of clades with preferences for colder or warmer regions. For example, Polygonaceae, Caryophyllaceae, and Montiaceae each are dominated by taxa with preferences for cold environments, although each also contains early-diverging taxa with preferences to warm environments. In contrast, taxa inhabiting warm environments predominate in Cactaceae, Amaranthaceae, Aizoaceae, the carnivorous clade (Droseraceae, Drosophyllaceae, Nepenthaceae, Ancistrocladaceae, Dioncophyllaceae), and the phytolaccoid clade (Nyctaginaceae, Phytolaccaceae, Petiveriaceae, Sarcobataceae, and *Agdestis*). Bioclimatic variable 12 (mean annual precipitation) showed a relatively consistent pattern of relatively dry to intermediately wet clades throughout the group. Indeed, only a few clades inhabiting wet ecosystems (in this case, the wet tropics) exist in the Caryophyllales, specifically small groups within the carnivorous clade, the phytolaccoids, early-diverging Polygonaceae, and other small groups throughout the Caryophyllales. The principal component loadings are presented in Fig. 2 and Fig. S5. Principal component 1, PCA1, showed significant differentiation throughout the Caryophyllales, as for example, early-diverging Polygonaceae vs the rest of Polygonaceae, early diverging Caryophyllaceae vs the rest of Caryophyllaceae, phytolaccoids vs Aizoaceae, and Portulacineae + relatives vs Cactaceae, to mention a few. These results generally reflect the extensive ecological diversification throughout the group. They also reflect significant diversification in the temperate regions of the world especially within the Caryophyllaceae and Polygonaceae contrasted with extensive diversification in the succulent lineages (especially Aizoaceae and Cactaceae) found in relatively dry and warm environments.

**Figure 1.**
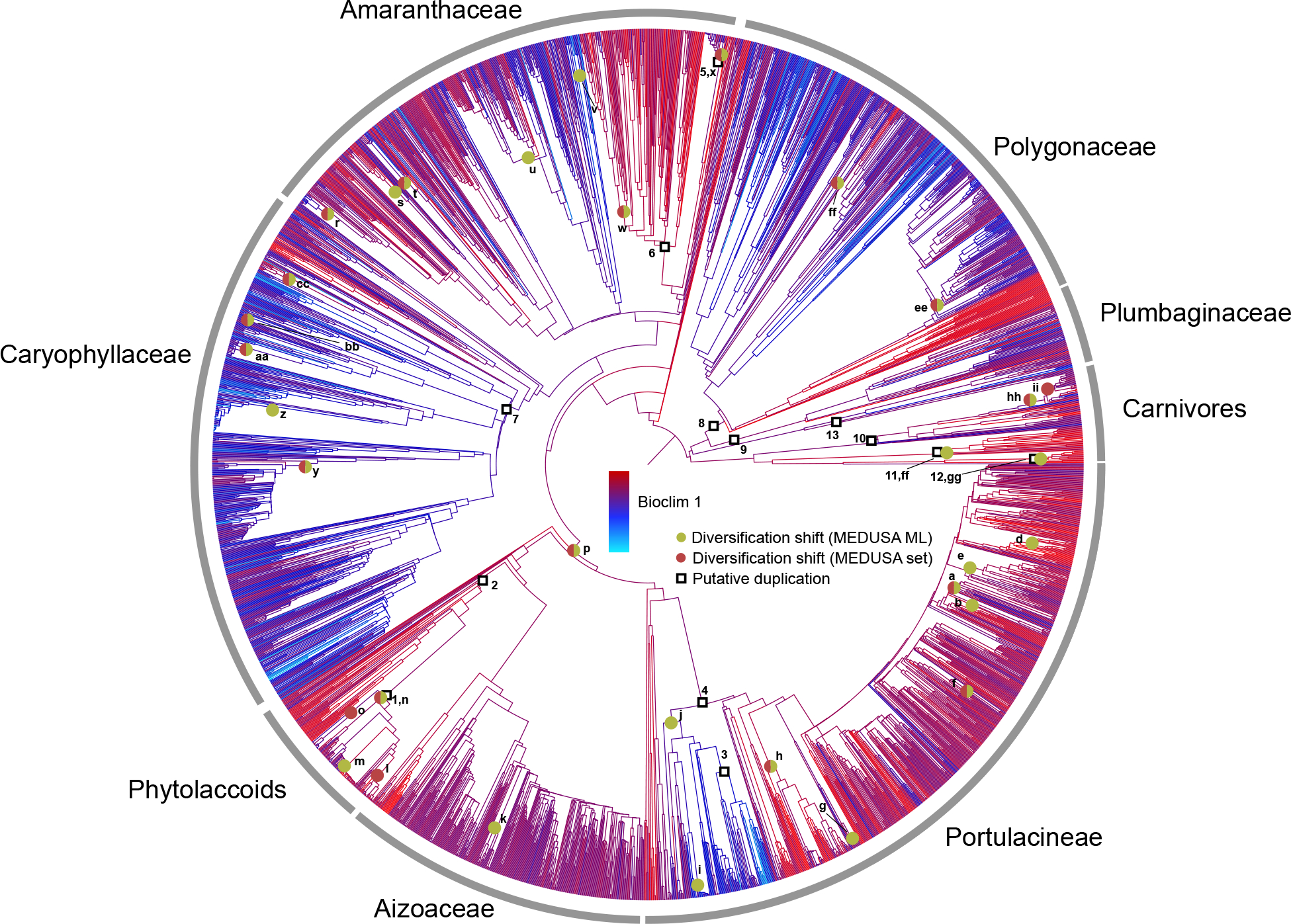
Chronogram of the Caryophyllales with putative WGD mapped along with identified diversification shifts. Diversification analyses were performed on the maximum likelihood tree (ML) as well as the bootstrap tree set (set) and those shifts that were identified in both groups are shown. The branches are colored based on Bioclim variable 1 (Mean Annual Temperature).

**Figure 2.**
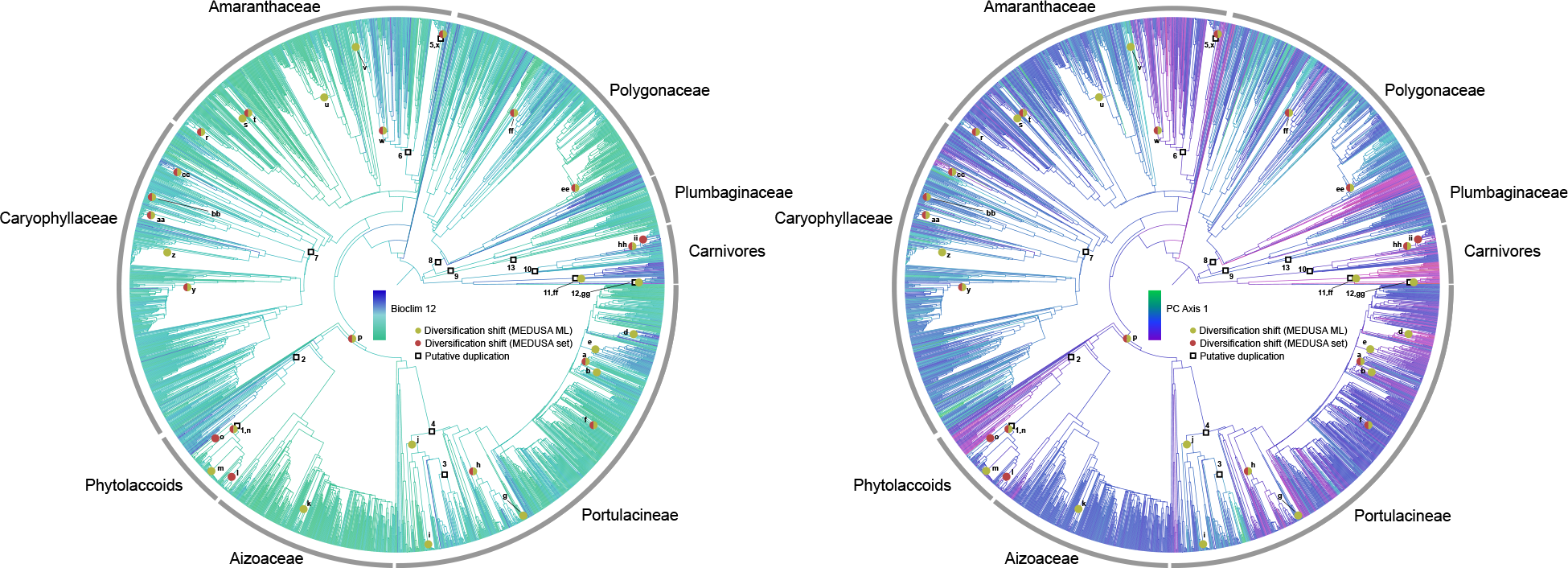
The chronograms and mapping of diversification and WGD are as in Fig. 1 (see caption for details). A) The branches are colored based on Bioclim variable 12 (Mean Annual Precipitation), and B) based on the principal component analyses (PCA) axis 1.

**Figure 3.**
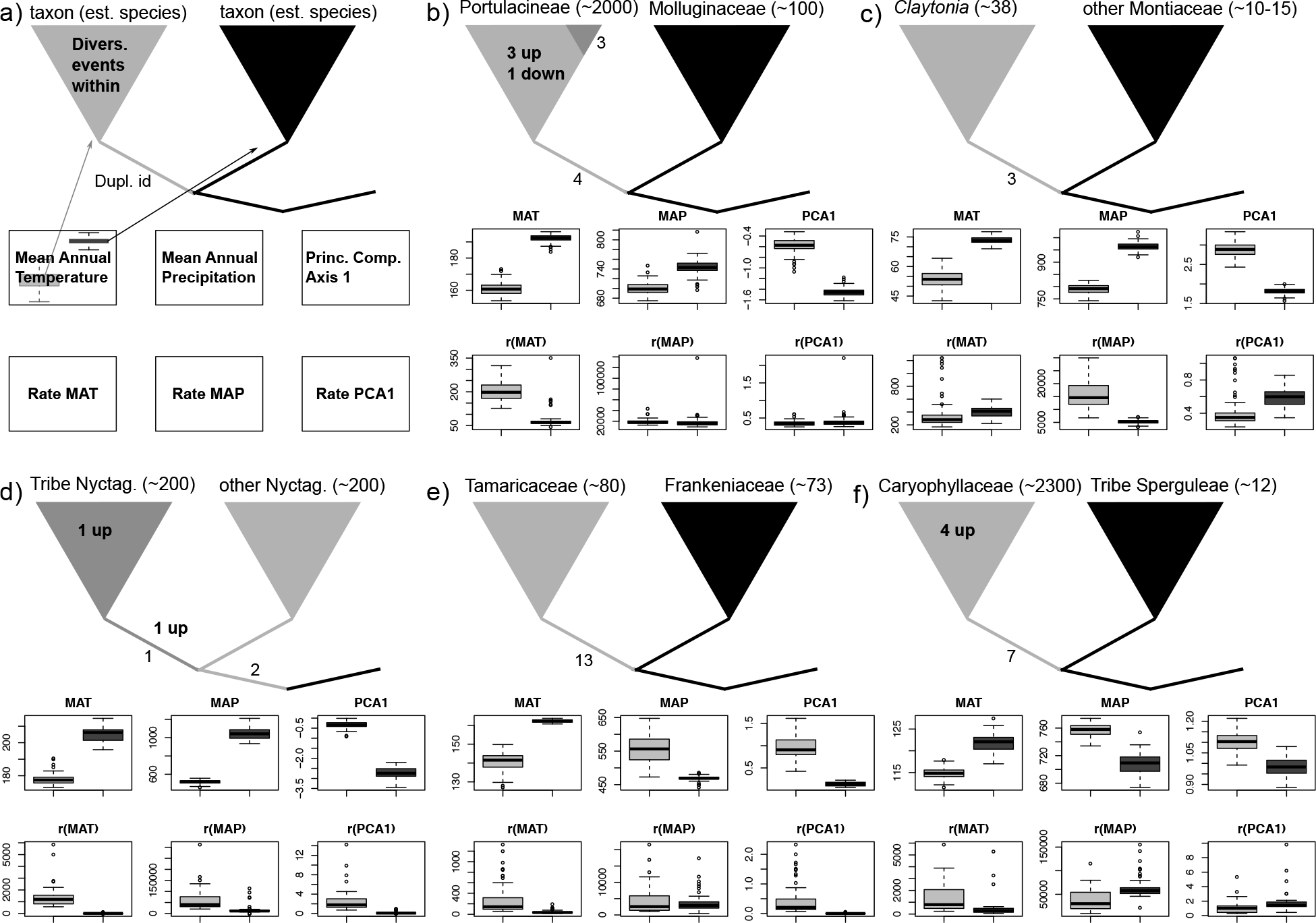
Summary of WGD events, with numbers corresponding to those in Table 3, climatic variables, and diversification shifts. Numbers along branches denote WGD, with the numbers corresponding to those in Fig. 1 and Table 3. Numbers inside clades denote the number of diversification rate shifts. Estimated species numbers are listed beside clade names. Box plots show the values estimated (ancestral values are listed in the top rows, rates in the bottom rows) for both the left and right clades across bootstrap samples. Clades shaded grey denote a WGD. b), c), and d) have nested WGD.

### Diversification

Significant shifts in diversification were detected in most major clades (Table 4, Fig. 1). The results from diversification analyses on the maximum likelihood tree and bootstrap tree set were generally congruent with each other. However, there were discrepancies (Fig. 1). The bootstrap set recovered many shifts in Polygonaceae, the carnivorous clade, Caryophyllaceae, some shifts within Cactaceae, phytolaccoids, and Amaranthaceae. Disagreements on the existence and placement of shifts are primarily within Portulacineae, Aizoaceae, and Amaranthaceae. Overall, MEDUSA detected 27 increases in diversification rate using the ML tree and 16 increases using the bootstrap trees. Given the relative lack of support of some of the branches in the phylogeny, we find the MEDUSA results on the set of bootstrapped trees to be the most conservative while the ML results are suggestive but not definitive of diversification shifts.

### Duplications, diversification, and climate occupancy

WGD analyses showed thirteen putative WGDs that can be mapped to clades (i.e., involve more than 1 taxon in the dataset; Table 3 and Figs. 1-3). Many of these were found in early diverging lineages as opposed to nested deep within families, though there are WGDs identified in *Amaranthus* and *Claytonia*. We also found evidence of nested WGDs as within the phytolaccoids and Portulacineae. In addition to these deeper WGDs, there were several more recent WGDs that were present in Ks plots but could not be mapped to a clade (Yang *et al*., submitted). By sampling more extensively, Yang *et al*. (submitted) and Walker *et al*. (2017) found additional WGD events within the Caryophyllales. We will surely find additional WGDs events in other lineages as more effort is placed on denser taxon sampling using genomes and transcriptomes. We did not explore WGDs that could only be mapped to terminal branches as we could not verify these phylogenetically. Further discussion of specific results related to the WGDs themselves can be found in Yang et al. (submitted) and Walker et al. (2017).

**Table 3:** Summary of WGD events at identified clades with distance to diversification shift and climate occupancy information. Numbers correspond to those in Figs. 1 and 2.

| # | Putative WGD | Distance to diversification shift in nodes ML(BS) | Subtending species (sister) | Mean annual temp °C (sister) | Mean annual precip mm (sister) |
| --- | --- | --- | --- | --- | --- |
| 1 | Tribe Nyctagineae within the Nyctaginaceae | 0 (0) | 123 (40) | 17.49 (20.08) | 482.9 (997.08) |
| 2 | Phytolaccoid clade | 6 (6) | 182 (407) | 19.64 (18.36) | 1007.58 (452.47) |
| 3 | *Claytonia* | NA | 38 (15) | 5.28 (7.25) | 790.5 (970.36) |
| 4 | Portulacineae | 1 (1) | 1600 (38) | 16.19 (19.35) | 699.87 (736.42) |
| 5 | *Amaranthus* | 0 (0) | 28 (1) | 16.27 (27.09) | 797.74 (117.63) |
| 6 | Tribe<br>Gomphrenoideae<br>within<br>Amaranthaceae | 7 (7) | 172 (41) | 17.91 (16.65) | 871.95 (1289.5) |
| 7 | in Caryophyllaceae<br>(Alsinoideae +<br>Caryophylloideae<br>sensu Greenberg and<br>Donoghue 2011) | 9 (9) | 793 (13) | 11.44 (12.06) | 761.43 (720.00) |
| 8 | Polygonaceae | 13 (13) | 670 (70) | 16.3. (16.89) | 1084.17<br>(794.28) |
| 9 | Plumbaginaceae | NA | 70 (670) | 16.89 (16.3) | 794.28<br>(1084.17) |
| 10 | Droseraceae | 8 (NA) | 67 (108) | 16.3 (19.08) | 1280.57<br>(1491.72) |
| 11 | Nepenthaceae | 4 (NA) | 89 (19) | 22.52 (20.05) | 2170.5<br>(1611.63) |
| 12 | Ancistrocladaceae | 0 (NA) | 15 (3) | 24.17 (25.6) | 1899.13<br>(2882.4) |
| 13 | Tamaricaceae | NA | 19 (3) | 14.09 (16.21) | 568.32 (469.61) |

**Table 4:** Summary of diversification shifts. Letters correspond to those in Figs. 1 and 2.

| # | Family | Diversification shift | Mean shift (ML) | Mean shift (BS) |
| --- | --- | --- | --- | --- |
| a | Cactaceae | *Echinops* | 1.7957 | 2.2008 |
| b | Cactaceae | within *Gymnocalycium* | 6.9152 |  |
| c | Cactaceae | *Gymnocalycium* | -0.001 | 0.0555 |
| d | Cactaceae | *Hylocereus*+*Selenicereus* | 0.1175 |  |
| e | Cactaceae | *Rhipsalis*+*Schlumbergera*+*Echinocereus*+relatives | 0.0514 |  |
| f | Cactaceae | *Stenocactus* | -0.057 | -0.019 |
| g | Anacampserotaceae | *Anacampseros* | 0.2624 |  |
| h | Portulacaceae | *Portulaca* | 0.0427 | 0.0447 |
| i | Montiaceae | *Montiopsis* | 0.9418 |  |
| j | Montiaceae | Montiaceae | 0.0325 |  |
| k | Aizoaceae | *Drosanthemum*+*Delosperma*+*Hereroa*+relatives | 0.1469 |  |
| l | Nyctaginaceae | *Boerhavia* |  | 0.0747 |
| m | Nyctaginaceae | *Commicarpus* | 0.9642 |  |
| n | Nyctaginaceae | Tribe Nyctagineae | 0.0484 | 0.0485 |
| o | Nyctaginaceae | *Abronia* |  | -0.084 |
| p | Nyctag.+Aizo+Cact.+relatives | Nyctag.+Aizo+Cact.+relatives | 0.0168 | 0.019 |
| r | Amaranthaceae | *Salicornia* | 0.2732 | 0.1649 |
| s | Amaranthaceae | *Suaeda* clade 1 | 0.1027 |  |
| t | Amaranthaceae | *Suaeda* clade 2 | -0.036 | -0.028 |
| u | Amaranthaceae | *Atriplex* | 0.0384 |  |
| v | Amaranthaceae | *Corispermum* | 0.1186 |  |
| w | Amaranthaceae | *Froelichia*+*Gomphrena*+relatives | 0.0217 | 0.0132 |
| x | Amaranthaceae | *Amaranthus* | 0.335 | 0.2049 |
| y | Caryophyllaceae | *Dianthus* | 0.0662 | 0.0409 |
| z | Caryophyllaceae | *Cerastium* | 0.7137 |  |
| aa | Caryophyllaceae | *Arenaria* | 0.4606 | 0.425 |
| bb | Caryophyllaceae | *Moehringia* | 1.0971 | 0.995 |
| cc | Caryophyllaceae | *Schiedea* | 0.2339 | 0.2767 |
| dd | Polygonaceae | *Fagopyrum* | -0.04 | -0.034 |
| ee | Polygonaceae | *Eriogonum*+relatives | 0.0432 | 0.0364 |
| ff | Nepenthaceae | within *Nepenthes* | 0.042 |  |
| gg | Ancistrocladaceae | *Ancistrocladus* | 0.1426 |  |
| hh | Droseraceae | within *Drosera* 1 | 0.2237 | 0.2076 |
| ii | Droseraceae | within *Drosera* 2 |  | 0.1622 |

To better examine whether WGDs coincide with diversification rate shifts, increases and decreases, or notable changes in climate tolerance, we mapped WGDs onto the large phylogenies and summarized the number of species and climate occupancy information for each clade (Tables 3-4, Figs. 1-3). Some WGD events were associated with synchronous diversification events. For example, within Nyctaginaceae, a WGD event occurs on the same branch (leading to Tribe Nyctagineae; Douglas & Spellenberg, 2010) as an increase in diversification rate in both the ML tree and the bootstrapped dataset (Fig. 1, dup:1 div:n). These events were also associated with a shift in life history and niche from an ancestral woody habit in the tropics to the largely herbaceous, arid-adapted temperate Nyctagineae. This was also the case for *Amaranthus* (Fig. 1, dup:5 div:x). Other coincident diversification and WGD events in the Droseraceae and Nepenthaceae were only supported by the ML tree. Although these correlated events may, in fact, be accurate, we will reserve more comments for when these are more confidently resolved. Other than these simultaneous shifts and one diversification shift at the base of the MRCA of Nyctaginaceae+Cactaceae, all other shifts in diversification occured more recently than WGD events. Several authors have suggested that this lagging pattern may be common at the broader angiosperm scale (Schranz *et al*. 2015, Tank *et al*. 2015), though the expected distance of the diversification shift from the WGD event was not specified (this is discussed more below). In the results presented here, some diversification events occur shortly after the WGD event, such as within the Amaranthaceae (dup: 6) and Portulacineae (dup: 4). For others, it is difficult to determine whether the diversification events that occur after the WGD events are significantly close to the WGD to warrant suggestion of an association (e.g., dup: 7, dup: 10, dup: 8). More description of a model that would generate a null expectation would be necessary to determine what is “close enough” (see discussion below).

Many of the other inferred lineage diversification rate shifts are associated with very recent, rapid radiations within genera such as those documented within *Commicarpus* (Nyctaginaceae), *Dianthus* (Caryophyllaceae), *Cerastium* (Caryophyllaceae), *Arenaria* (Caryophyllaceae), and *Salicornia* (Amaranthaceae), to name a few (Table 4). Although polyploids were reported in these clades, we were unable to pinpoint the phylogenetic location of any WGD with our current taxon sampling (e.g., *Dianthus*; Carolin, 1954; Weiss *et al*. 2002). Increased sampling of transcriptomes and genomes will shed more light in these areas. While we only find a few WGDs that coincide well with diversification rate shifts, it is important to note that the uncertainty in the phylogenies makes it difficult to map anything but the strongest diversification signals. This discrepancy can be seen in the difference between the number of events supported by the ML analyses and those supported by the bootstrap analyses. It is possible that additional sequence data will improve phylogenetic resolution and confidence, and that consequently additional diversification events will emerge.

Equally interesting to the few WGD events associated directly with diversification are the WGD events associated with general shifts in climate tolerance. WGDs in the Polygonaceae, Caryophyllaceae, Montiaceae, and the Tribe Nyctagineae appear to be associated with movement into colder environments (Figs. 1-2 and Figs. S2-S3). Species arising after the WGD within the Amaranthaceae occupy wetter environments than the sister clade. The WGDs within the carnivorous plants were also associated with shifts in environment as Nepenthaceae are found in very wet environments and the Droseraceae are found in somewhat drier environments, at least comparatively. However, in these cases, perhaps the development of the wide array of morphologies associated with carnivory, apart from *Drosophyllum*, is more obviously associated with the WGD (Walker *et al*., 2017).

While these qualitative assessments suggest potential correlations of shift in the climate occupied and WGDs, more specific and direct comparisons are necessary to quantify the extent of the shifts. For many of the clades experiencing WGD, a direct comparison with a sister clade is difficult because the sister may consist of a single species, another clade with WGD, or another complication. For example, there are WGDs at the base of both Polygonaceae and Plumbaginaceae as well as Nepenthaceae and Droseraceae. However, we made direct comparison of five duplicated lineages (see Fig. 3) in both means (i.e., character contrasts between sister clades) and variances (rate of Brownian motion) of climatic variables. In each case, the duplicated lineage occupied a colder mean annual temperature. This was also the case with the nested WGDs of Portulacineae and the Tribe Nyctagineae. Of course, we are not suggesting that all WGDs are associated with a shift to a colder climate. While such a pattern may exist in some groups such as Caryophyllaceae, we emphasize the observation that there is a shift in the climate occupied rather than the direction of the shift. Mean annual precipitation is not as clear with some clades occupying a higher precipitation and some occupying lower precipitation. Perhaps the best summary of climatic niche is the principal components of all the climatic variables. Here, while the shift in units is less easily interpreted, duplicated clades occupied different niches than sister lineages. This supports the hypothesis that WGD events are associated with adaptations. Here, many of these adaptations are associated with shifts in climatic niches. This necessitates further examination in other angiosperm clades to investigate how general these results are.

The rates of niche evolution show more complicated patterns. While some clades, such as the Portulacineae, showed significant increase in a rate of niche evolution as compared to the sister clade (e.g., MAT), no clear pattern emerged across all comparisons. There were other shifts in rate such as with MAT and MAP in the Nyctaginaceae and Montiaceae, but these were not as strong as the pattern of climate occupancy itself discussed above.

With each of these patterns presented here, it is important to consider them in the context of uncertainty, both inherent in the biological processes that generate the phylogeny and in the analyses associated with large scale datasets. These large phylogenies and datasets allow for more thorough examination of the clades, but uncertainty makes precise mapping of weaker signals difficult. As mentioned above, both the mapping of diversification events and duplications demonstrate this. Furthermore, the comparisons of the sister clades for climatic niche analyses assumes accurate identification of sister lineages. Increasing taxon sampling may help, but additional sequence data and specimen data for phylogenetic analyses, WGD mapping analyses, and climate niche characterization will surely improve our precision in these investigations.

What emerges from these analyses of WGD, diversification, and climate occupancy? It would appear as though, perhaps not unexpectedly, the patterns are complex and mixed. Some WGD are associated directly with diversification events, some WGD are associated with shifts in climate tolerance, some WGD are coincident with shifts in rates of niche evolution, and still other WGD are associated with known adaptations (carnivory, habit shifts associated with montane habitats, etc.). Some diversification shifts follow WGD events. However, it is unclear whether these events are linked or correlated and, if so, if they are correlated more with diversification than an additional adaptation or other evolutionary pattern or process. As data increase in these groups and as confidence increases in the phylogenetic relationships as well as the placement of both diversification and WGD events, we will be able to better address these questions. However, at least for the Caryophyllales, it does not appear as though diversification is tightly linked with WGD. Instead, for the clades that can be tested, we find shifts in climate occupancy correspond well to WGD.

### Suggestions for moving forward

WGD are almost certainly one of the dominant processes that contribute to major evolutionary events within plant lineages. This may be in the form of increased diversification, development of novel traits, adaptation to new environments, and many other events (e.g., Schubert and Vu, 2016; Clavijo *et al*. 2017). However, for several reasons, these events (i.e., WGD and other evolutionary events) may not occur simultaneously. In fact, there may be little to no expectation for the events to occur simultaneously (e.g., Donoghue, 2005; Schranz *et al*. 2012; Donoghue & Sanderson, 2015; Tank *et al*., 2015; Dodsworth *et al*. 2016). In any case, more precise expectations and null models need to be developed to allow for reasonable tests of the correlations among these events. For example, there may be shifts in diversification that follow a WGD, but is it close enough, or frequent enough to infer that the two events are related? Is correlation possible or identifiable if, as is expected, intervening lineages have gone extinct? These questions would benefit from simulation studies where the true correlation pattern is known. Furthermore, more precise connections should be made to the biology of speciation and genome WGDs to better determine why, specifically, WGDs would be expected to correspond with any diversification pattern instead of adaptations, which may or may not correspond with increases or decreases in speciation. While still challenging, investigating the fate of and patterns of selection within individual genes (e.g., subfunctionalization and neofunctionalization) may shed light into the genomic basis of post-WGD and possibly allow for more concrete expectations for diversification. With the availability of genomes and transcriptomes, this is now beginning to become a possibility (e.g., Brockington *et al*., 2015, Walker *et al*., 2017). Only when these suggestions are linked to more specific biological hypotheses will we be able to better understand the ultimate impact of WGD in plant evolution.

## Acknowledgements

We thank Caroline Parins-Fukuchi for discussion of the project and comments on the manuscript. We thank Gregory Stull, Oscar Vargas, Ning Wang, Sonia Ahluwalia, Jordan Shore, Lijun Zhao, Alex Taylorm, and Drew Larson for helpful comments on the manuscript. The authors thank Hilda Flores, Helga Ochoterena, Tom Wendt and the staff at the Plant Resources Center at the University of Texas at Austin, the Lyon Arboretum, David Anderson, John Brittnacher, Anna Brunner, Joseph Charboneau, Arianna Goodman, Heather-Rose Kates, Patricia Herrnández Ledesma, Lucas Majure, Nidia Mendoza, Michael Powell, Rick Ree, Carl Rothfels, Flora Samis, Jeffrey Sanders, Elizabeth Saunders, Rich Spellenberg, Greg Stull, Mats Thulin, Erin Tripp, and Sophia Weinmann for help with obtaining material. We thank the Cambridge University Botanic Gardens for growing material for this study. This work was supported by NSF DEB awards 1352907 and 1354048.

## Author contributions

S.A.S., J.F.W., Y.Y., M.J.M., and S.F.B. designed research. C.P.D, R. B., N.L., and N.A.D. collected data. S.A.S., J.W.B., and Y.Y. analyzed the data. S.A.S. led the writing. All authors read and contributed to the manuscript.

## Supporting Information

**Fig.S1** The cladogram with support mapped for the bootstrap replicates described in the methods.

**Fig.S2** The chronograms and mapping of temperature variables (bioclimatic variables 13-19) that are not presented in Fig. 1.

**Fig.S3** The chronograms and mapping of precipitation variables (bioclimatic variables 13-19) that are not presented in Fig. 2.

**Fig.S4** The chronograms and mapping of PCA axis 2 on the broader Caryophyllales.

**Fig.S5** Principal component loadings for bioclimatic variables.

